# Tanshinone IIA potentiates the therapeutic efficacy of glucocorticoid in lipopolysaccharide-treated HEI-OC1 cells through modulation of Foxp3/Nrf2 signaling pathway

**DOI:** 10.1101/2024.08.19.608552

**Authors:** Jie Li, Xiaoyan Zhu, Shiming Ye, Qi Dong, Jie Hou, Jing Liu, Wandong She

## Abstract

Glucocorticoids (GC) are commonly used to treat sudden sensorineural hearing loss (SSNHL), although some patients show resistance to this therapeutic approach. Clinical studies demonstrate the efficacy of tanshinone IIA (TA) in combination with GC for managing various human ailments. However, it remains unclear whether TA can mitigate GC resistance in SSNHL.

**Aim of the study:** Our aim is to elucidate the role of NRF2-induced transcriptional regulation of HDAC2 in influencing GC resistance and investigate the involvement of TA-related molecular pathways in GC resistance.

**Materials and Methods:** HEI-OC1 cells are treated with lipopolysaccharide (LPS) to establish an in vitro model for SSNHL. Subsequently, the cells are treated with dexamethasone (DXE) or DXE+TA. RT-qPCR and western blot analyses are employed to measure mRNA and protein levels of Forkhead box P3 (FOXP3), nuclear factor erythroid 2-related factor 2 (NRF2), and histone deacetylase 2 (HDAC2). Cell Counting Kit-8 (CCK-8) and 5-ethynyl-2’-deoxyuridine (EdU) assays are conducted to assess cell proliferation. Flow cytometry analysis is performed for apoptosis evaluation. Mechanistic studies involve Chromatin immunoprecipitation (ChIP), luciferase reporter, and DNA pull-down assays.

**Results:** Treatment with TA+DEX significantly enhances proliferation and suppresses apoptosis in LPS-treated HEI OC1 cells. TA upregulates HDAC2 expression by activating NRF2-mediated transcription of HDAC2, with the NRF2-HDAC2 binding site located at bases 419-429 (ATGACACTCCA) in the promoter sequence of HDAC2. Furthermore, TA upregulates FOXP3 expression to activate NRF2 transcription, with the predicted FOXP3-binding site located at bases 864-870 (GCAAACA) in the promoter sequence of NRF2.

**Conclusion:** This study’s findings suggest that TA enhances the therapeutic effects of GC on proliferation and apoptosis in HEI OC1 cells by up-regulating FOXP3/Nrf2 expression. These results indicate that TA may be promising in ameliorating GC resistance in patients with SSNHL.

## Introduction

Sudden sensorineural hearing loss (SSHL) is a prevalent otological emergency. Studies have shown that SSHL affects 5 to 27 per 100,000 people per year, with nearly 66,000 new cases annually in the United States [1]. Sensory hearing loss often occurs due to damaged or deficient cochlear hair cells; hair cells may be abnormal at birth or damaged during an individual’s lifetime. Proposed etiologies of primary SSHL include virus infection, vascular insufficiency, autoimmune disorders, and stress theory [2,3]. There is no proven or recommended treatment or cure for SSHL. The treatment of SSHL remains one of the most challenging issues in contemporary otorhinolaryngology. Therefore, it is necessary to explore promising therapeutic strategies for patients with SSHL. **HEI-OC1 is one of the few mouse auditory cell lines available for research. Originally proposed as an in vitro system for screening of ototoxic drugs, these cells have been used to investigate drug-activated apoptotic pathways, autophagy, senescence, mechanism of cell protection, inflammatory responses, cell differentiation, genetic and epigenetic effects of pharmacological drugs, effects of hypoxia, oxidative and endoplasmic reticulum stress, and expression of molecular channels and receptors** [4]**. Currently, the HEI-OC1 cell line has been widely used in the field of hair cell research. To a certain extent, it can be used to speculate on in vivo changes and establish a hair cell injury and inflammation model.**

Currently, glucocorticoids (GCs) are widely used to treat SSHL and various immune and inflammatory diseases [5–8].Although most patients with SSHL respond well to GCs, about 20% of patients show no significant hearing improvement after GC treatment, indicating GC resistance in these patients [9,10].

Fortunately, traditional Chinese medicine has been combined with Western medicine to prevent and reverse drug resistance in cancer cells [11]. This approach has also been applied to the treatment of SSHL. For instance, treating SSHL patients with a combination of GC and breviscapine (a traditional Chinese medicine) has been proven more effective than using GC alone [12]. Therefore, it is worthwhile to explore the potential of traditional Chinese medicine in preventing and reversing GC resistance in SSHL.

Tanshinone IIA (TA) is a bioactive compound derived from the rhizomes and roots of Salvia miltiorrhiza Bunge, commonly known as Danshen, a traditional Chinese herb. TA has garnered attention for its therapeutic potential in various diseases, including liver disorders [13], atherosclerosis, cancers, and other diseases [14–16]. At the molecular level, TA influences various pathways. For instance, it attenuates neuroinflammation by suppressing the RAGE/NF-κB signaling pathway, thereby potentially impeding the progression of Alzheimer’s disease [17]. Moreover, TA protects hair cells against radiation-induced damage by suppressing p65/NF-κB nuclear translocation and modulating the p53/p21 signaling pathway [18]. Furthermore, TA sodium sulfonate injection has shown promise in enhancing hearing and improving the hemorheological parameters and immune functions of SSHL patients. However, the molecular mechanisms underlying TA in SSHL remain to be explored.

Recent research has proposed a correlation between diminished nuclear factor erythroid 2-related factor 2 (NRF2) levels and histone deacetylase 2 (HDAC2) proteins and GC resistance in SSHL patients [10,19]. Our previous studies revealed decreased expression of HDAC2 and the apoptosis-inhibiting genes Bcl-2 and Bcl-xl in an LPS-induced hearing loss animal model and an LPS-treated HEI-OC1 cell injury model [20]. Furthermore, elevated NRF2 expression has mitigated GC insensitivity by promoting HDAC2 levels *in vitro* [21,22].

Forkhead box P3 (FOXP3) is a pivotal transcriptional factor regulating the expression of its target RNAs [23]. However, its involvement in SSHL remains largely unexplored. We applied bioinformatics tools to analyze the relationship between FOXP3 and NRF2 in the HEI-OC1 cell injury model. We focused on the role of NRF2-mediated transcriptional modulation of HDAC2 to elucidate the impact of TA-related molecular pathways on GC resistance. Our experimental results show that the combination of TA and DEX can significantly improve LPS-induced hair cell damage, which provides a new therapeutic strategy for the clinical treatment of SSHL.

## Materials and methods

### Cell culture and treatment

The HEI-OC1 cell line obtained from Professor Chai Renjie’s Laboratory (Southeast University Life Science Research Institute, Nanjing, China) was maintained in DMEM supplemented with 10% fetal bovine serum (FBS) and 100 U/mL of penicillin. Another cell line, 293T cells, purchased from ATCC, was maintained in DMEM with 10% FBS and 2 mM of glutamine.

As previously described, the HEI-OC1 cell injury model was induced by treating with 0.1 μg/mL of LPS (MCE, NJ, USA) for 24h [24]. To detect the effect of dexamethasone (DEX) on LPS-induced apoptosis in HEI-OC1 cells, 50 μg/mL of dexamethasone (DEX) was used to treat the cells as previously described [25]. TA purchased from MCE was used to treat the cells along with DEX 10 μM TA, a concentration that could significantly inhibit LPS-induced inflammation as previously described [26], was used in the current study.

### Plasmid transfection

Three small interfering RNAs (siRNAs) targeting NRF2 (si-NRF2-1, si-NRF2-2, si-NRF2-3) or HDAC2 (si-HDAC2-1, si-HDAC2-2, si-HDAC2-3) were synthesized by GenePharma (Shanghai, China) for silencing of each target, respectively. Non-targeted random siRNA was used as the negative control (si-NC). For over-expression of NRF2 or FOXP3, the full length of each target gene sequence was separately sub-cloned into a pcDNA3.1 expression vector while the pcDNA3.1 empty vector was used as the control group. Plasmids were transfected into HEI-OC1 cells using Lipofectamine 2000 (Invitrogen, Carlsbad, CA, USA). Cells were harvested for subsequent experiments after they were cultured for 48 hours.

### Real-time quantitative polymerase chain reaction (RT-qPCR)

Total RNA was isolated using TRIzol reagent (Sangon, Shanghai, China). The integrity and purity of the extracted total RNA were measured using a NanoDrop One ultra-micro-UV spectrophotometer (Thermo Fisher Scientific, Waltham, USA). A total of 2 μg RNA was used for reverse transcription. M-MLV Reverse Transcriptase (Sigma-Aldrich, St. Louis, MO, USA) was used to reverse transcription isolated RNA into cDNA. Reverse transcription was conducted at 25□°C for 5 min, 42□°C for 60 min, and 70□°C for 5 min. PCR was then performed using SYBR Green PCR Master Mix (Takara, Kyoto, Japan) for 40 cycles (95°C for 3 sec and 60°C for 30 sec). Relative RNA level was calculated with the 2^-ΔΔCt^ method by normalizing to the internal control (GAPDH).

### Western blot

Total protein was extracted from HEI-OC1 cells using RIPA lysis buffer. The supernatant was collected after centrifuging at 10,000 g for 10 min at 4 °C. The BCA protein assay (Pierce, Rockford, IL, USA) determined protein concentration. Proteins were separated on 10% SDS-PAGE gels and transferred onto PVDF membranes. The membranes were blocked with 5% non-fat milk in TBST for 1 h, then incubated with one of the primary antibodies: anti-BCL-2 (1: 2000, Proteintech Group, Inc., Rosemont, IL, USA), anti-BAX (1:2000, Proteintech Group, Inc.), anti-GR (1:20000, Proteintech Group, Inc.), anti-NRF2 (1:1000 dilution, Proteintech Group, Inc.), anti-HDAC2 (1:20000, Proteintech Group, Inc.), anti-FOXP3 (1:1000 dilution, Proteintech Group, Inc.) or anti-β-actin (1: 20000, Proteintech Group, Inc.) at 4°C overnight. After being washed four times with TBST, the membranes were incubated with a secondary antibody at room temperature for 2□h. Protein blots were visualized on an enhanced chemiluminescence (ECL) system (Tanon, Shanghai, China). β-actin was used as an internal reference. Image-Pro Plus 6.0 was used for density quantization of all protein bands.

### Cell Counting Kit-8 (CCK-8) assay

For CCK-8 assay, stably transfected cells were seeded into 96-well plates at 3000 cells per well. After 24 h incubation for attachment, the cells were incubated with fresh medium with LPS, DEX, or DEX+TA for 0, 24, 48, or 72 h. Subsequently, the cells were incubated with 10 μL of CCK-8 solution at 37□°C for another 2□h. The absorbance value at 450 nm was measured by an EnVision® multimode plate reader (Perkin Elemer).

### 5-ethynyl-2’-deoxyuridine (EdU) assay

Cell proliferation was assessed using an EdU assay kit (Ribobio, Guangzhou, China). In brief, transfected cells were seeded into 96-well plates and incubated with EdU at 37°C for 2 h. The cells were washed in PBS and fixed in 4% paraformaldehyde (Sangon Biotech Co., Ltd.) at 4°C for 30 min. Nuclei were stained with 400 μl of DAPI (Sigma-Aldrich; Merck KGaA) at room temperature for 30 min. Cell proliferation was observed using a Leica fluorescence microscope (x200). The percentage of EdU-positive cells in five randomly selected fields was calculated using Image J software v1.8.0 (National Institutes of Health, Bethesda, MD, USA).

### Apoptosis assay

According to the instructions, cell apoptosis was detected using an Annexin-V-FITC and Propidium Iodide (PI) Apoptosis Detection Kit (BD Biosciences, CA, USA). Briefly, cells were collected and washed twice in cold PBS. Cells were then fixed with 4% paraformaldehyde (Sangon Biotech Co., Ltd.) at −20°C overnight, followed by suspending in 600 μl of eBioscience™ flow cytometry binding buffer (Invitrogen; Thermo Fisher Scientific, Inc.) at a concentration of 106 cells/ml, and then stained with 5 μl of Annexin V/FITC and 5 μl of propidium iodide (BD Biosciences) in the dark for 15 min. Flow cytometry assessed apoptosis cells in triplicates (FACS Calibur, Becton Dickinson, NJ, USA). Results were analyzed using FlowJo 7.6 software (Tree Star, Inc., USA).

### Chromatin immunoprecipitation (ChIP) assay

Magna ChIP™ RNA-Binding Protein Immunoprecipitation Kit (Millipore, Billerica, MA, USA) was used for ChIP assay. Briefly, cells were cross-linked with 1% formaldehyde for 10 min and sonicated to shear DNA into 200∼1000 base pairs. Cell lysates were incubated with protein A/G beads coated with anti-NRF2 or anti-FOXP3 (Proteintech) at 4□°C overnight. Anti-rabbit IgG was used as a negative control. After washing, the bead-bound immunocomplexes were eluted using an elution buffer. To de-crosslink the DNA-protein complex, samples were added with 5 M of NaCl, heated at 65□°C for 4□h, then treated with proteinase K, and further incubated at 45□°C for 1□h. Finally, the immunoprecipitated DNA was purified for RT-qPCR analyses.

### Luciferase reporter assay

The whole sequence of HDAC2 promoter with or without HDAC2 [named wild type (WT) and mutant type (MUT)] was inserted into pGL3 luciferase vector to generate pGL3-HDAC2 promoter-WT or pGL3-HDAC2 promoter-MUT. Similarly, the whole sequence of NRF2 promoter with or without FOXP3 was inserted into pGL3 luciferase vector to generate pGL3-NRF2 promoter-WT or pGL3-NRF2 promoter-MUT. To demonstrate the functional sites, the pGL3-HDAC2 promoter-WT and pGL3-HDAC2 promoter-MUT were co-transfected with pcDNA3.1-NRF2 or pcDNA3.1 empty vector into 293T cells. Similarly, pGL3-NRF2 promoter-WT and pGL3-NRF2 promoter-MUT were co-transfected with pcDNA3.1-FOXP3 or pcDNA3.1 empty vector into 293T cells. Transfections were completed using Lipofectamine 2000 Transcription Reagent (Invitrogen; Thermo Fisher Scientific, Inc.) at 37°C for 46 h. Finally, the luciferase activity of Firefly was detected using a Luciferase Reporter Assay System (Promega) by normalizing it to Renilla luciferase activity.

### DNA pull-down assay

Biotinylated wild-type or mutated HDAC2 promoter/NRF2 promoter probes (Bio-HDAC2 promoter-WT/MUT or Bio-NRF2 promoter-WT/MUT) or NC probes (Bio-NC) synthesized by Ribobio were added to 1% Triton X-100, followed by heating at 95 °C for 2 min and cooling in an ice bath for 3 min. The denatured biotinylated RNA was mixed with 15 µl of streptavidin beads (Thermo Fisher Scientific, Inc.) to pull down the biotinylated protein-DNA complex. The DNA-bead complexes were incubated with 100 ml of cell lysate obtained using RIPA lysis buffer. Finally, proteins eluted from DNA were subjected to western blot.

### Statistical analysis

Data from three independent experiments were shown as mean ± standard derivation (SD) by plotting them using GraphPad Prism v8.0 (San Diego, CA, USA). SPSS V22.0 software (IBM, Armonk, NY, USA) was used for statistical analysis of all data. The difference between the two groups was compared by using an unpaired Student’s t-test. The difference among multiple groups was compared using a one-way analysis of variance (ANOVA) with Dunnett or Tukey post hoc text. P<0.05 indicated that the difference was statistically significant.

## Results

### TA enhances HDAC2 protein level in LPS-treated HEI-OC1 cells

In the present study, we utilized an in vitro LPS injury model [20]to investigate the impact of TA on GC sensitivity in SSHL. We conducted proliferation and apoptosis assays on HEI-OC1 cells treated with 0.1 μg/mL of LPS for 24-72 hours. Cell proliferation assessed via CCK-8 and EdU assays was markedly suppressed by LPS treatment compared to the control group (P < 0.01, Figure 1A, B). Meanwhile, LPS treatment markedly increased the apoptotic rate (P < 0.01, Figure 1C), which was further proved by the elevated BAX protein levels and reduced BCL-2 protein levels (Figure 1D). Additionally, we evaluated HDAC2 protein expression in HEI-OC1 cells treated with LPS, LPS+DEX, or LPS+DEX+TA. HDAC2 protein levels were reduced in HEI-OC1 cells treated with LPS alone (P < 0.01, Figure 1E), while this reduction was partially reversed by DEX treatment and further restored by combined treatment with DEX and TA (P < 0.01, Figure 1E).

**Figure 1.**
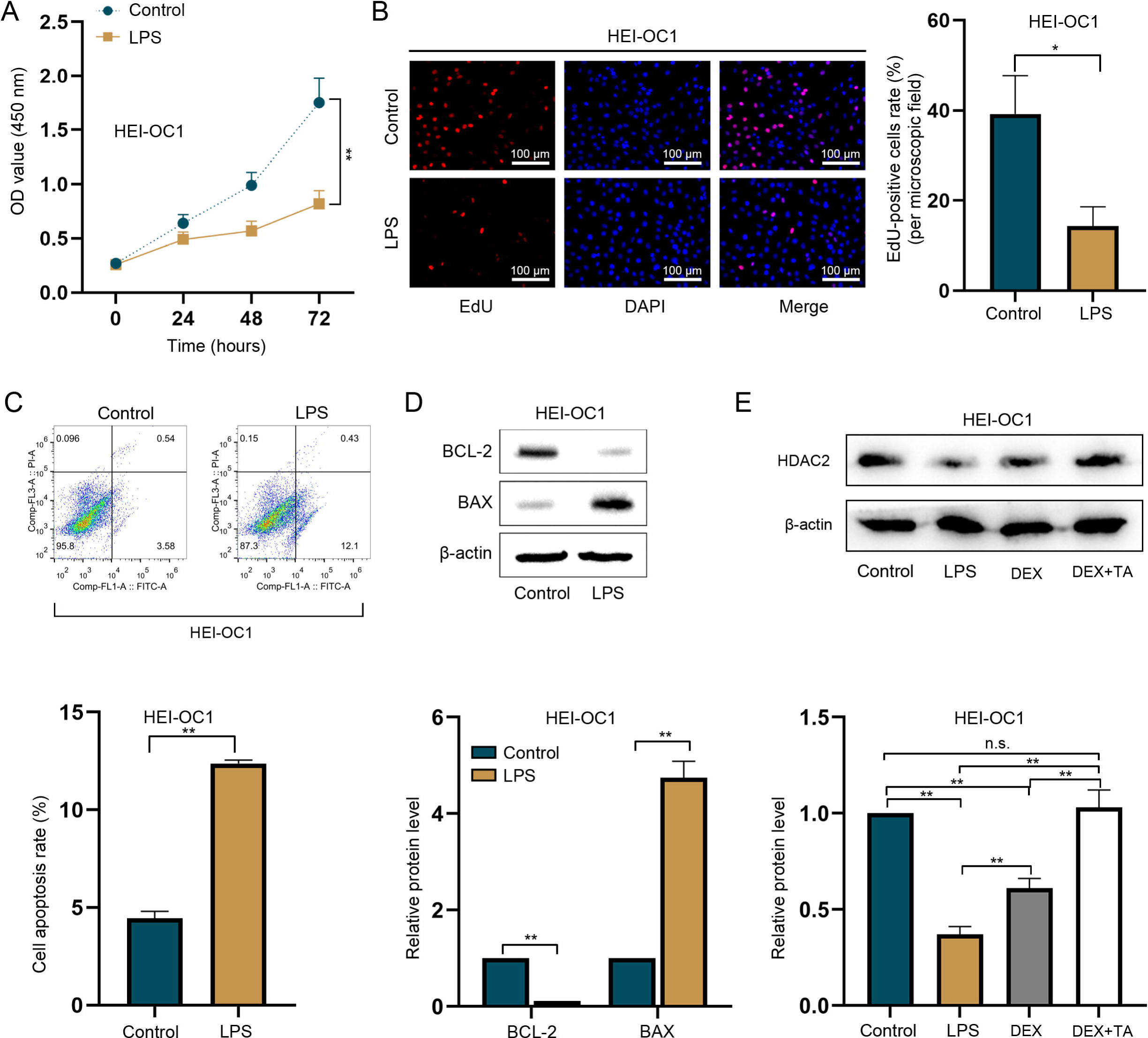
TA enhances HDAC2 protein level in HEI-OC1 cells treated with LPS. A-B. CCK-8 and EdU assays were used to detect proliferation in HEI-OC1 cells treated with LPS. As detected by CCK-8 and EdU assays, cell proliferation was markedly suppressed by LPS treatment compared to the control group. Scale bar = 100 µm in B. C. Apoptosis in HEI-OC1 cells treated with LPS was assessed via flow cytometry analysis. The apoptotic rate was increased significantly by LPS treatment. D. Western blot analysis of BCL-2 and BAX expressions in HEI-OC1 cells treated with LPS. The BAX protein level increased while the BCL-2 protein level decreased after LPS treatment. E. Western blot analysis of HDAC2 protein expression in HEI-OC1 cells treated with LPS, DEX, or DEX+TA. HDAC2 protein levels were decreased in HEI-OC1 cells treated with LPS alone, while its levels were partly reversed by DEX treatment and totally reversed by treatment with both DEX and TA (**P < 0.01).

### TA enhances proliferation and GR expression and inhibits apoptosis in LPS-treated HEI-OC1 cells

We then evaluated the effects of TA on proliferation and apoptosis in LPS-treated HEI-OC1 cells. Treated 50 μg/mL of DEX significantly increased the proliferation rate in HEI-OC1 cells (P < 0.05), while combined treatment of DEX (50 μg/mL) +TA (10 μM) treatment induced a higher proliferation rate in HEI-OC1 cells (P < 0.01, Figure 2A-B). Moreover, DEX inhibited LPS-induced apoptosis of HEI-OC1 cells (P < 0.01), and additional TA further suppressed apoptosis more effectively than DEX alone (P < 0.01, Figure 2C). Western blots further supported these findings, showing that DEX increased BCL-2 protein levels and decreased Bax protein levels, with TA significantly enhancing the anti-apoptotic effects of DEX (Figure 2D).

**Figure 2.**
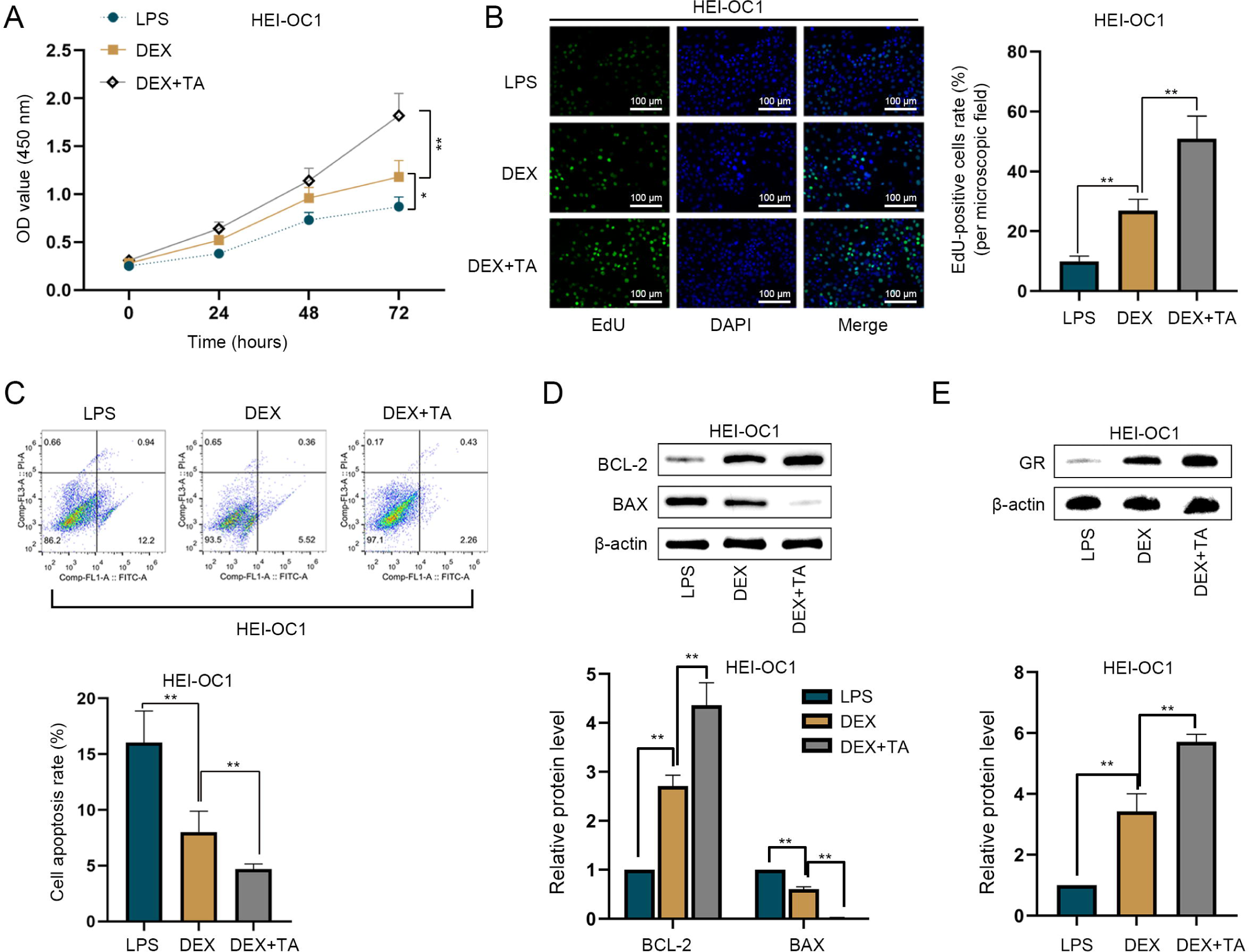
TA increases GR level and proliferation and inhibits apoptosis in HEI OC1 cells treated with LPS. A-B. CCK-8 and EdU assays were used to detect proliferation in LPS-treated HEI-OC1 cells. It turned out that 50 μg/ mL of DEX treatment increased the proliferation rate in HEI-OC1 cells treated with LPS, while DEX+TA (10 μM) treatment induced a higher proliferation rate in HEI-OC1 cells treated with LPS. Scale bar: 100 µm. C. Apoptosis induced by LPS in HEI-OC1 cells was assessed via flow cytometry analysis. DEX inhibited apoptosis induced by LPS in HEI-OC1 cells (p), and the additional TA treatment intensively suppressed apoptosis compared to DEX alone (p). D. Western blot analysis of BCL-2 and BAX expressions in HEI-OC1 cells treated with LPS. DEX alone increased BCL-2 protein level and decreased Bax protein level (p), while TA significantly enhanced the inhibitory effect of DEX on cell apoptosis (p). E. Western blot analysis of GR expression in HEI-OC1 cells. GR protein level was elevated after TA treatment compared with DEX treatment alone in HEI-OC1 cells treated with LPS (*P < 0.05 and **P < 0.01).

Previous work has revealed that GR mediates the physiologic and pharmacologic actions of GCs [9]. Consequently, we investigated the impact of TA on GR expression. Western blot demonstrated that the GR protein level was elevated after TA treatment compared with DEX treatment alone in LPS-treated HEI-OC1 cells (P < 0.01, Figure 2E). Previous studies suggest that increased GR and HDAC2 levels and enhanced cell proliferation are associated with reduced GC resistance [6,9,10]. Therefore, TA can potentially inhibit GC resistance by upregulating GR and HDAC2 expression, promoting proliferation, and inhibiting apoptosis.

### TA enhances HDAC2 expression through NRF2-mediated transcription activation

Recent research has demonstrated low NRF2 and HDAC2 expression levels in patients with refractory SSHL [10]. Our previous study also indicated that reduced NRF2, HDAC2, and GR expression levels are associated with decreased GC sensitivity in SSHL patients [9,10,19]. HDAC2 is a critical component of the GC-GR complex, which mediates the trans-repression of NF-кB transcriptional activity by deacetylating histones in the pro-inflammatory genes and deacetylating GR [9]. We hypothesized that TA might regulate NRF2 and HDAC2 expression, thereby influencing GC sensitivity in SSHL patients.

We observed that TA further increased NRF2 and HDAC2 mRNA and protein levels in DEX-treated HEI-OC1 cells (P < 0.01, Figure 3A), suggesting that TA could reverse GC resistance in SSHL via upregulating NRF2 and HDAC2 expression. Then, we explored the potential regulatory relationship between NRF2 and HDAC2. After confirming the high efficiency of NRF2 overexpressing (P < 0.01, Figure 3B), we detected increased HDAC2 mRNA levels in HEI-OC1 cells overexpressing NRF2 (P < 0.01, Figure 3C). Given the role of NRF2-mediated transcriptional regulation [27], we speculated that NRF2 might activate HDAC2 transcription to up-regulate its expression.

**Figure 3.**
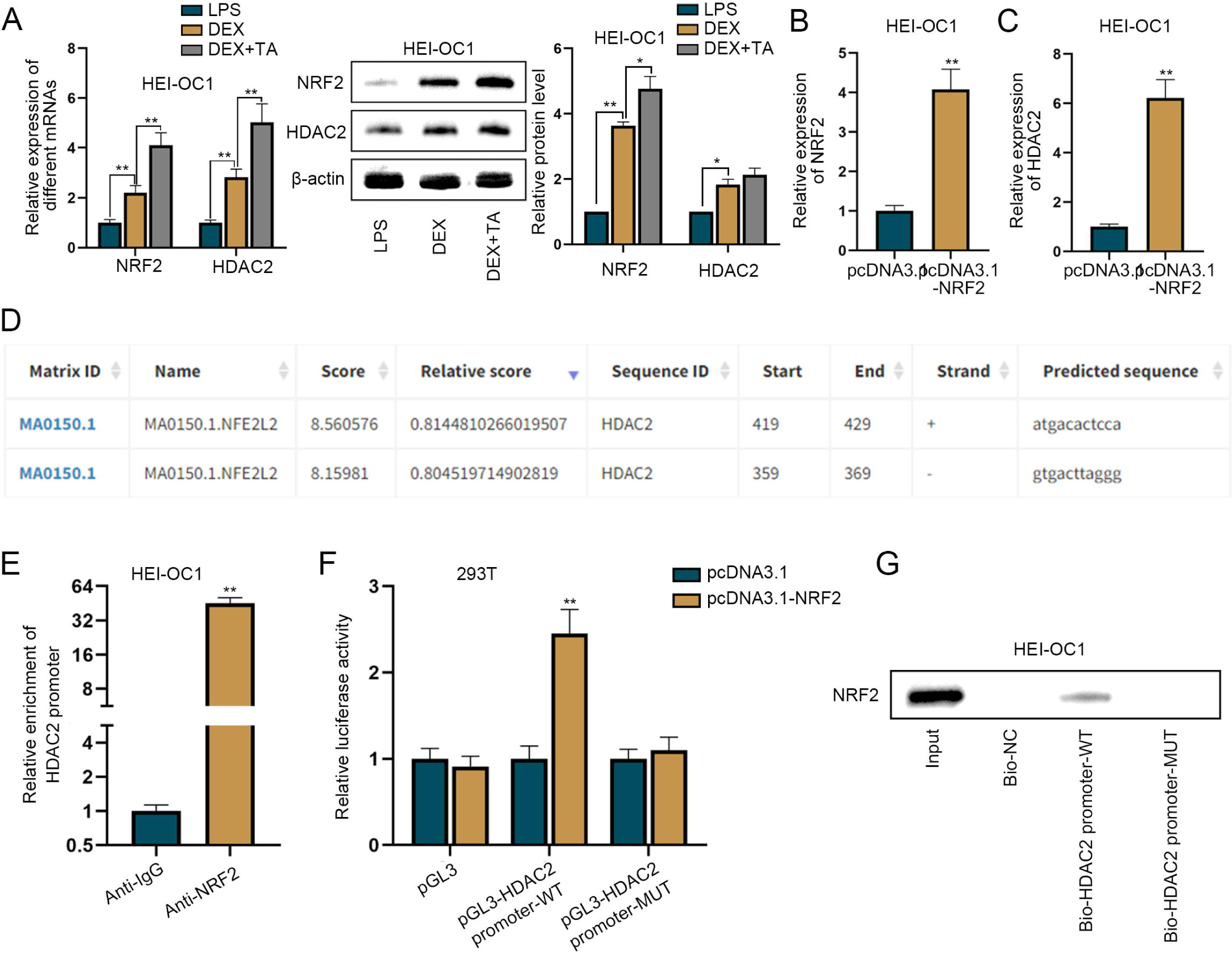
TA enhances HDAC2 expression through NRF2-mediated transcription activation. A. q-PCR and western blot analyses of NRF2 and HDAC2 expressions in HEI-OC1 cells treated with LPS, DEX or DEX+TA. TA increased NRF2 and HDAC2 mRNA and protein levels in DEX-treated HEI-OC1 cells (**P < 0.01). B. RT-qPCR was used to check gene over-expression of NRF2 in HEI-OC1 cells. NRF2 was successfully over-expressed in HEI-OC1 cells. C. HDAC2 mRNA level was detected via RT-qPCR in HEI-OC1 cells over-expressed NRF2. We detected an increased HDAC2 mRNA level in HEI-OC1 cells over-expressed NRF2 (**P < 0.01). D. Potential NRF2-binding site in the sequence of HDAC2 promoter was predicted by JASPAR database. Based on JASPAR database (https://jaspar.genereg.net/) analysis, we obtained putative NRF2-binding sites in the sequence of HDAC2 promoter. The NRF2-binding site with the highest prediction score is located at 419–429 bases (ATGACACTCCA) in HDAC2 promoter sequence. E. ChIP detected enrichment of HDAC2 promoter in immunoprecipitate with Anti-NRF2. ChIP assay presented an abundant enrichment of HDAC2 promoter in immunoprecipitates with Anti-NRF2, verifying the binding between NRF2 and HDAC2 promoter. F. Luciferase activity of different reporter constructs in NRF2-overexpressing 293T cells was detected. G. DNA pull-down assay followed by western blot was used to detect NRF2 expression in the complex pulled down by Bio-HDAC2 promoter-WT. NRF2 over-expression increased the luciferase activity of HDAC2 promoter-WT while not changing the HDAC2 promoter-MUT activity (mutated sequence: TACTGTGAGGT at 419–429 bases). **P < 0.01.

Based on the JASPAR database (https://jaspar.genereg.net/), we identified putative NRF2-binding sites in the HDAC2 promoter sequence, with the highest prediction score located at base 419–429 (ATGACACTCCA) (Figure 3D). ChIP assay results showed significant enrichment of HDAC2 promoter in immunoprecipitates with anti-NRF2 (Figure 3E), verifying the binding between NRF2 and HDAC2 promoter. A Luciferase reporter assay indicated that NRF2 overexpression increased the luciferase activity of HDAC2 promoter-WT but did not affect the activity of the HDAC2 promoter-MUT (mutated sequence: TACTGTGAGGT at 419–429 bases, Figure 3F). DNA pull-down assay also illustrated that NRF2 directly binds to HDAC2 promoter-WT rather than HDAC2 promoter-MUT (Figure 3G). Taken together, TA upregulates HDAC2 expression by activating NRF2-mediated HDAC2 transcription.

### TA up-regulates FOXP3 expression to activate NRF2 transcription

Subsequently, we investigated the underlying mechanism in which TA enhances NRF2 expression. Previous studies have shown that TA can enhance FOXP3 expression to inhibit nasopharyngeal carcinoma progression [28]. Measuring FOXP3 expression in LPS-treated HEI-OC1 cells, we found that FOXP3 mRNA and protein levels were significantly increased in response to DEX treatment and further elevated with additional TA treatment (all P < 0.01, Figure 4A). Transcription factor FOXP3 is expressed in various tissues and is crucial in anti-tumor and anti-inflammatory activities [29]. Hence, we speculated that TA up-regulates NRF2 expression might be via FOXP3.

**Figure 4.**
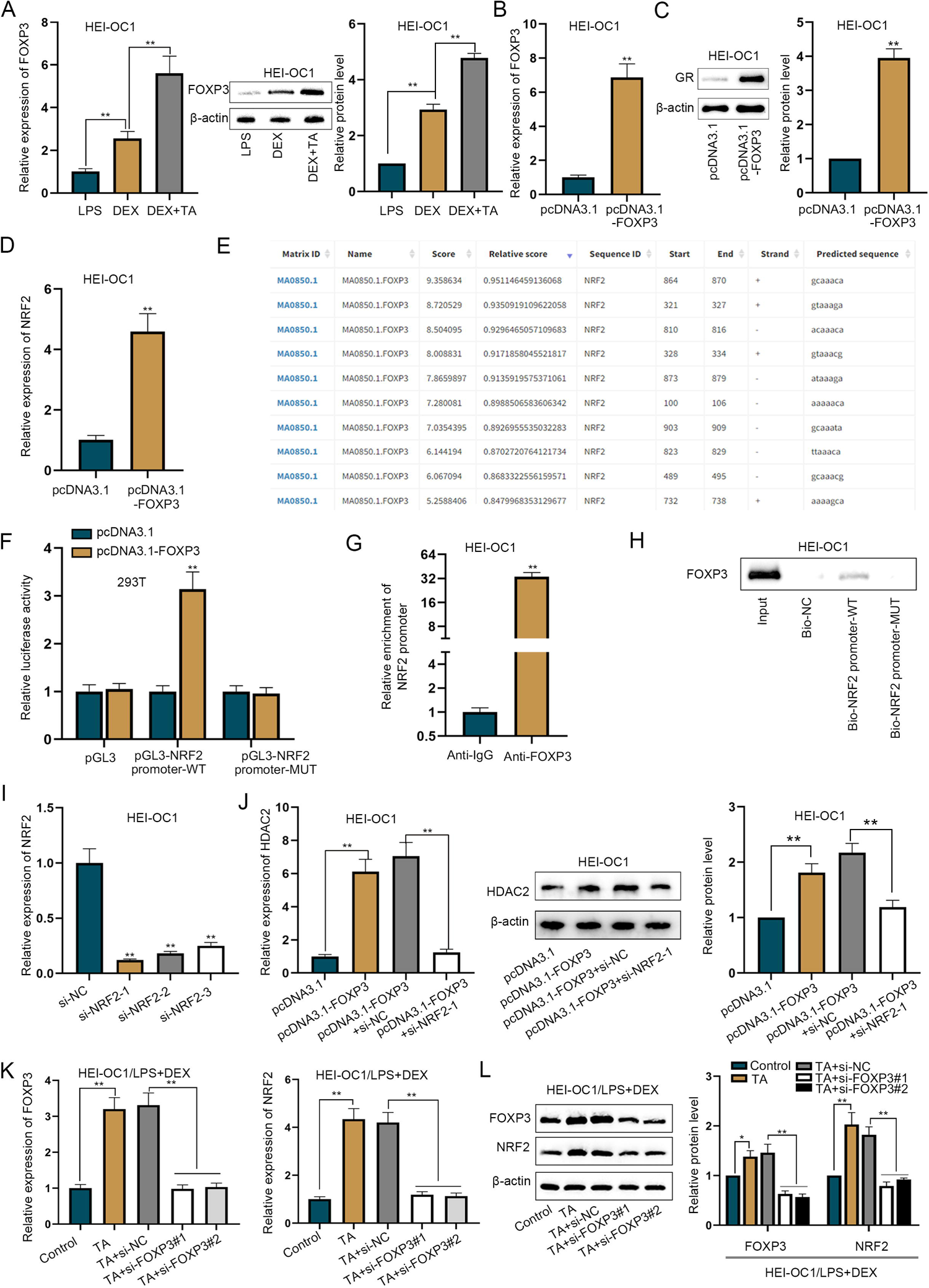
TA up-regulates FOXP3 expression to activate NRF2 transcription. A. RT-qPCR and western blot were used to measure FOXP3 expression in LPS-treated HEI-OC1 cells. FOXP3 mRNA and protein levels were increased in response to additional TA treatment (P < 0.01). B. Gene over-expression efficiency of FOXP3 in HEI-OC1 cells was tested via RT-qPCR. FOXP3 was over-expressed in HEI-OC1 cells. C. GR expression in FOXP3-over-expressing HEI-OC1 cells was measured via western blot. GR protein level was also increased in FOXP3-over-expressing HEI-OC1 cells. D. NRF2 expression in FOXP3-over-expressing HEI-OC1 cells was measured via RT-qPCR. Elevated NRF2 mRNA level was observed in FOXP3-over-expressed HEI-OC1 cells. E. Potential FOXP3-binding sites in NRF2 promoter sequence were predicted by JASPAR database. According to JASPAR database analysis, the predicted FOXP3-binding site with the highest prediction score is located at 864–870 bases (GCAAACA) in NRF2 promoter sequence. F. Luciferase activity of different reporter constructs in FOXP3-over-expressing 293T cells was detected by luciferase reporter assay. The luciferase activity of pGL3-NRF2 promoter-WT was enhanced by up-regulation of FOXP3 in 293T cells, while that of pGL3-NRF2 promoter-MUT (mutated sequence: CGTTTGT at 864–870 bases) was not affected. G. ChIP assay was performed to detect the physical binding between FOXP3 and NRF2 promoter. ChIP assay uncovered that NRF2 promoter was highly enriched in Anti-FOXP3-precipitated complex. H. DNA pull-down assay followed by western blot was used to detect FOXP3 expression in the complex pulled down by Bio-NRF2 promoter-WT. DNA pull-down assay validated the effectiveness of the binding sites between FOXP3 and NRF2 promoter. I. The knock-down efficiency of si-NRF2 plasmids was checked via RT-qPCR. All three siRNAs had high efficacy of NRF2 knock-down. J. HDAC2 expression was measured via RT-qPCR and western blot in HEI-OC1 cells. The increased mRNA and protein levels of HDAC2 caused by FOXP3 over-expression were reversed by depletion of NRF2. K. FOXP3 expression was knocked down in DEX-treated HEI OC1 cells. The mRNA level of NRF2 was measured in DEX-treated HEI OC1 cells after FOXP3 knock-down by RT-qPCR. L. Protein levels of FOXP3 and NRF2 were measured in DEX-treated HEI OC1 cells after FOXP3 knock-down. The mRNA and protein levels of NRF2 increased by TA treatment while the levels were re-decreased by silencing of FOXP3 (**P < 0.01).

Upon successful overexpressing FOXP3 in HEI-OC1 cells (P < 0.01, Figure 4B), we observed an increased GR protein level was also increased (P < 0.01, Figure 4C), and elevated NRF2 mRNA level (P < 0.01, Figure 4D). According to JASPAR database analysis, the predicted FOXP3-binding site with the highest prediction score is located at bases 864–870 (GCAAACA) in the NRF2 promoter sequence (Figure 4E). Luciferase reporter assay showed the luciferase activity of pGL3-NRF2 promoter-WT was enhanced by FOXP3 upregulation in 293T cells, while the activity of pGL3-NRF2 promoter-MUT (mutated sequence: CGTTTGT at 864–870 bases) was not affected (Figure 4F). ChIP assay result indicated that the NRF2 promoter was highly enriched in the anti-FOXP3-precipitated complex (Figure 4G). The DNA pull-down assay also validated the binding sites between FOXP3 and NRF2 promoter (Figure 4H). Further, we performed rescue experiments to analyze the regulatory relationship among FOXP3, HDAC2, and NRF2. First, we confirmed the high efficacy of NRF2 knockdown (Figure 4I) and found the increased mRNA and protein levels of HDAC2 caused by FOXP3 overexpression were reversed by NRF2 deletion (Figure 4J). To validate the effect of TA on the FOXP3/NRF2 axis, we knocked down FOXP3 expression with specific siRNAs in DEX-treated HEI-OC1 cells (Figure 4K, left panel) and examined NRF2 mRNA and protein levels. RT-qPCR and western blot showed TA treatment increased NRF2 mRNA and protein levels, which were decreased again by silencing FOXP3 (Figure 4K, right panel & Figure 4L).In summary, TA increases FOXP3 expression, which activates NRF2-mediated HDAC2 transcription, to up-regulate HDAC2 expression.

### FOXP3 alleviates LPS-induced cell apoptosis via HDAC2

A previous study indicated that TA increases FOXP3 expression to inhibit the progress of nasopharyngeal carcinoma [28]. We found that FOXP3 mRNA and protein levels were significantly elevated in HEI-OC1 cells treated with LPS, followed by DEX and TA (P < 0.01,Figure 4A). To verify the regulatory effect of the FOXP3/HDAC2 axis in LPS-treated HEI-OC1 cells,we conducted rescue experiments.

Firstly, we knocked down HDAC2 by transfecting si-HDAC2. Among the siRNA tested, si-HDAC2-1 plasmid showed the highest interference efficiency and was used in subsequent experiments (Figure 5A). HDAC2 knock-down with si-HDAC2-1 abrogated the proliferative effect of FOXP3 overexpression on HEI-OC1 cells (Figure 5B-C). The elevated BCL-2 protein levels and decreased BAX protein levels resulting from FOXP3 overexpression were reversed by HDAC2 deficiency (Figure 5D). Moreover, the reduction in the cell apoptosis rate caused by FOXP3 overexpression was restored by HDAC2 knock-down (Figure 5E). In summary, FOXP3 over-expression impedes LPS-induced apoptosis in HEI-OC1 cells by upregulating HDAC2.

**Figure 5.**
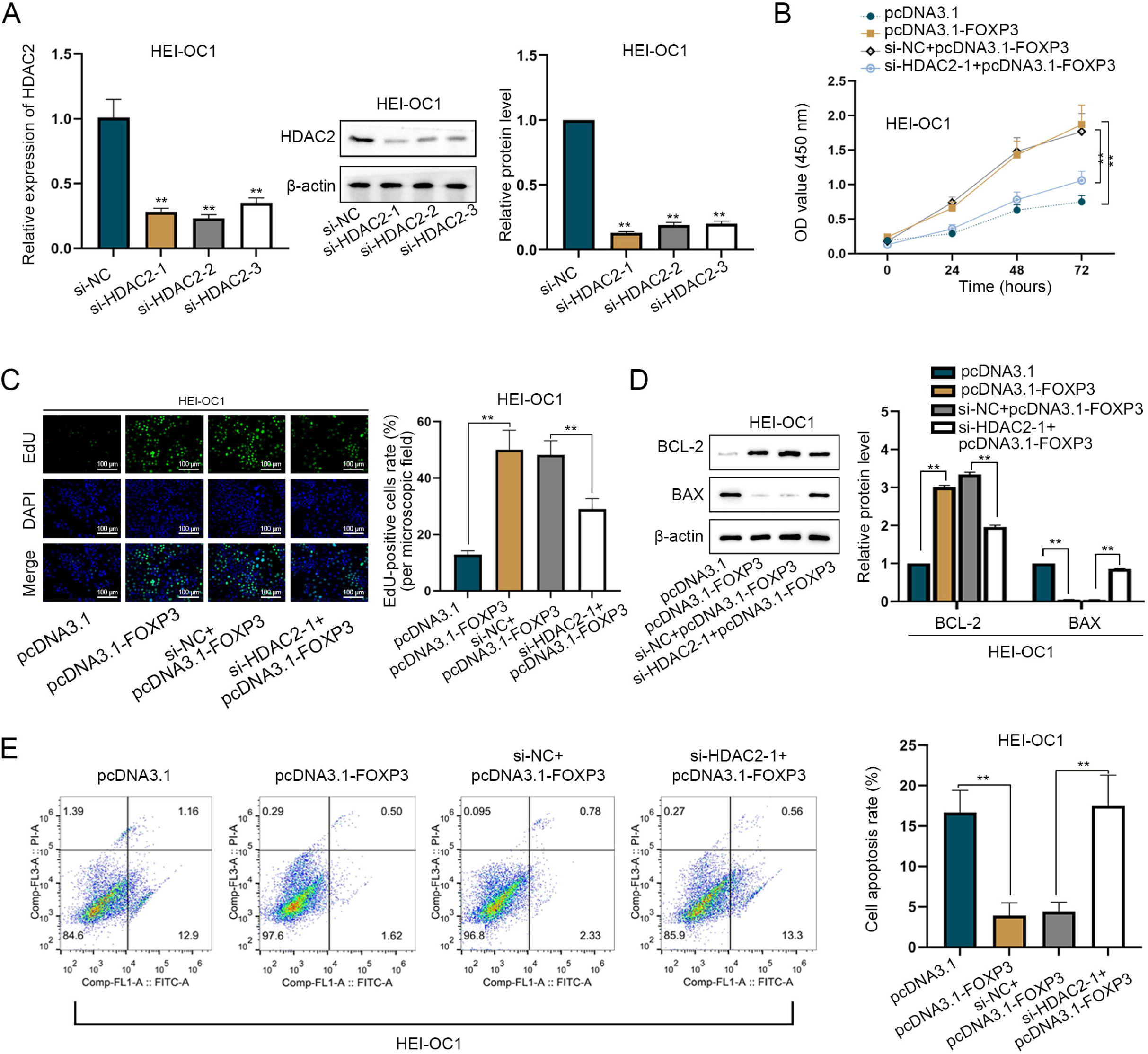
FOXP3 alleviates LPS-induced apoptosis via HDAC2. A. Interference efficiency of si-HDAC2 plasmids in LPS-induced HEI-OC1 cells was checked via RT-qPCR and western blot. si-HDAC2-1 plasmid was used in the subsequent experiments for its highest interference efficiency. B-C. CCK-8 and EdU assays were performed to assess cell proliferation. Scale bar: 100 µm. HDAC2 knock-down abrogated the promoting effect of FOXP3 over-expression on cell proliferation. D-E. Western blot of BAX and BCL-2 expressions and flow cytometry analysis were performed to evaluate apoptosis. Elevated BCL-2 protein levels and declined BAX protein levels resulting from FOXP3 up-regulation were recovered by HDAC2 deficiency. Moreover, the apoptosis rate reduced by FOXP3 over-expression was restored by HDAC2 knock-down (**P < 0.01).

## Discussion

SSHL is a prevalent emergency in otolaryngologic clinics. While GC was the first-line treatment for SSHL worldwide, resistance to GC in many SSHL patients remains a clinical challenge [30]. Recent studies have found that the expression of HDAC2 can enhance the sensitivity of GC treatment by affecting the deacetylation of glucocorticoid receptors. Reduced HDAC2 expression is considered to be an important cause of GC resistance in SSNHL [31]. HDAC2 activity is reduced through oxidative stress-induced post-transcriptional modifications, such as nitration of tyrosine residues and phosphorylation of serine residues, leading to resistance to GC in SSNHL patients [32]. To address drug resistance, the combination of traditional Chinese medicine combined with Western medicine has been explored. For example, a combination therapy of GC and breviscapine has shown to be more effective than GC alone for SSHL patients [12].

Previous reports have demonstrated the therapeutic value of TA in treating various diseases. Subedi and Gaire suggested TA as an appealing drug candidate for neurodegenerative diseases [33]. Xue et al. have manifested that TA could repress the proliferation and migration of human colon cancer cells [34]. Although TA exhibits anti-tumor activity by inducing autophagy and apoptosis and inhibits cell growth and migration by activating AMPK and inhibiting PI3K/Akt/mTOR signaling pathways, TA attenuates cell death in LPS- and hydrogen peroxide-induced inflammation models through NF-kB pathway [35]. The different effects of it in different models may be caused by different cell types, drug concentrations, and molecular targets of the drug. In the current research, we explored the effect of TA on proliferation and apoptosis in cochlear cells treated with LPS as an *in vitro* SSHL model. Consistent with our previous studies [20], LPS inhibited proliferation and increased apoptosis in HEI-OC1 cells. DEX treatment significantly promoted proliferation and inhibited apoptosis induced by LPS. Additional TA treatment significantly enhanced the effects of DEX on proliferation and apoptosis. Our molecular mechanism study indicates that TA upregulates HDAC2 expression by activating NRF2-mediated transcription of HDAC2 through FOXP3. The predicted NRF2 binding site in the HDAC2 promoter sequence is located at base 419–429 bases (ATGACACTCCA).

HDAC2 is a critical regulator of the cell cycle, apoptosis, proliferation, migration, and differentiation [36]. Recent advancements have highlighted the role of HDAC2 in SSHL. For instance, Hou et al. have proposed that reduced HDAC2 protein levels may be critical for corticosteroid insensitivity in patients with refractory SSHL [19]. Additionally, NRF2 is a transcription factor involved in mitochondrial biogenesis, autophagy, metabolic reprogramming, immunity, and inflammation [37]. Low expression levels of NRF2/HDAC2 proteins are linked to GC insensitivity in SSHL [10,19]. Compared with previous studies, our current study further investigated the mechanisms underlying the interaction of TA with NRF2/HDAC2 in an *in vitro* SSHL model. This study demonstrated that TA upregulates HDAC2 expression through the activation of NRF2-mediated HDAC2 transcription. Additionally, we validated that TA upregulates FOXP3 expression, which in turn activates NRF2 transcription. The predicted FOXP3-binding site is located at base 864–870 bases (GCAAACA) in the NRF2 promoter sequence.

The results of our *in vitro* experiments demonstrated that TA upregulates FOXP3 expression to activate the transcription of NRF2. NRF2 in turn promotes HDAC2 transcription, leading to enhanced HDAC2 expression. Elevated HDAC2 levels assist the transcriptional regulatory effect of the GR-GC complex, then increase cells’ response to GC [9,19]. This molecular mechanism suggests that TA could be a potential therapeutic agent to overcome GC resistance in SSHL patients.

However, our current results are based on in vitro experiments using HEI-OC1 cells. In future studies, the protective effect of TA+DEX combination therapy on SSHL should be further verified in mouse models and zebrafish models to provide more comprehensive and reliable experimental support for clinical treatment. The effects of TA treatment should be further validated in vivo studies before it can be considered for use in SSHL patients.

## Conclusion

The current study demonstrates that TA can synergistically enhance the treatment effects of GC on proliferation and apoptosis in HEI-OC1 cells through NRF2, HDAC2 and FOXP3, indicating that TA may have a therapeutic potential to ameliorate GC resistance in SSHL patients.

## Authorship contribution statement

Wandong She contributed to the study’s conception and design, funding, and critically revised the manuscript. Jie Li performed the experiment, data curation, formal analysis, writing, and editing the manuscript. Xiaoyan Zhu provided data curation and technical support and reviewed the manuscript. Shiming Ye and Qi Dong provided experimental technical support. Jie Hou and Jing Liu provided data curation and technical support. All authors read and approved the submitted version.

## Acknowledgments

We want to give our sincere thanks to Drs. Xiaoping Du and Zachary Yokell (Hough Ear Institute, Oklahoma, USA) thank you for their critical review and thoughtful feedback while preparing this manuscript.

## Abbreviation

SSNHL: sudden sensorineural hearing loss
GC: glucocorticoids
TA: tanshinone IIA
LPS: lipopolysaccharide
DXE: dexamethasone
FOXP3: Forkhead box P3
NRF2: nuclear factor erythroid 2-related factor 2
HDAC2: histone deacetylase 2
CCK-8: Cell Counting Kit-8
EdU: 5-ethynyl-2’-deoxyuridine
ChIP: Chromatin immunoprecipitation

